# Steviol Glycoside biosynthesis pathway gene expression profiling of transformed and non-transformed plant leaf tissues of *Stevia rebaudiana* (Bertoni)

**DOI:** 10.1101/2025.08.14.670226

**Authors:** Priya Singh, Amol Phule, Heena Tabassum, Minal Wani

## Abstract

*Stevia rebaudiana* Bertoni, a member of the Asteraceae family, is recognized for its sweetened leaves which are remarkably sweeter than sucrose by 200–300 times. This astounding property is because of steviol glycosides (SGs), a class of diterpenoid secondary metabolites which primarily consists of stevioside and rebaudioside A. These compounds are formed through a specific steviol glycoside (SG) biosynthetic pathway that contains several key genes. In the present work, gene expression profiling of 15 core genes of SG biosynthetic pathway, along with metabolite analysis were conducted in three groups of *Stevia rebaudiana* plants: *In vitro* regenerated non-transformed plantlets (NP), *in vitro* regenerated transformed plantlets (TP) via hairy root cultures using *Rhizobium rhizogenes* mediated transformation and Control plants (CP). Quantitative real-time PCR (qRT–PCR) results showed that in NP and TP there was upregulation of 13 genes. Both NP and TP showed downregulated *SrDXR* and *SrCDPS* in comparison to CP, whereas *SrUGT74G1* had higher expression in NP than TP. HPLC chromatographic studies on SGs showed that stevioside content followed the order TP > NP > CP. These findings demonstrate that transformation enhances SG biosynthesis and support the use of genetically modified *Stevia rebaudiana* lines for increased natural sweetener production. Further studies are warranted to elucidate regulatory mechanisms and optimize metabolic engineering approaches.

**Graphical Abstract:** 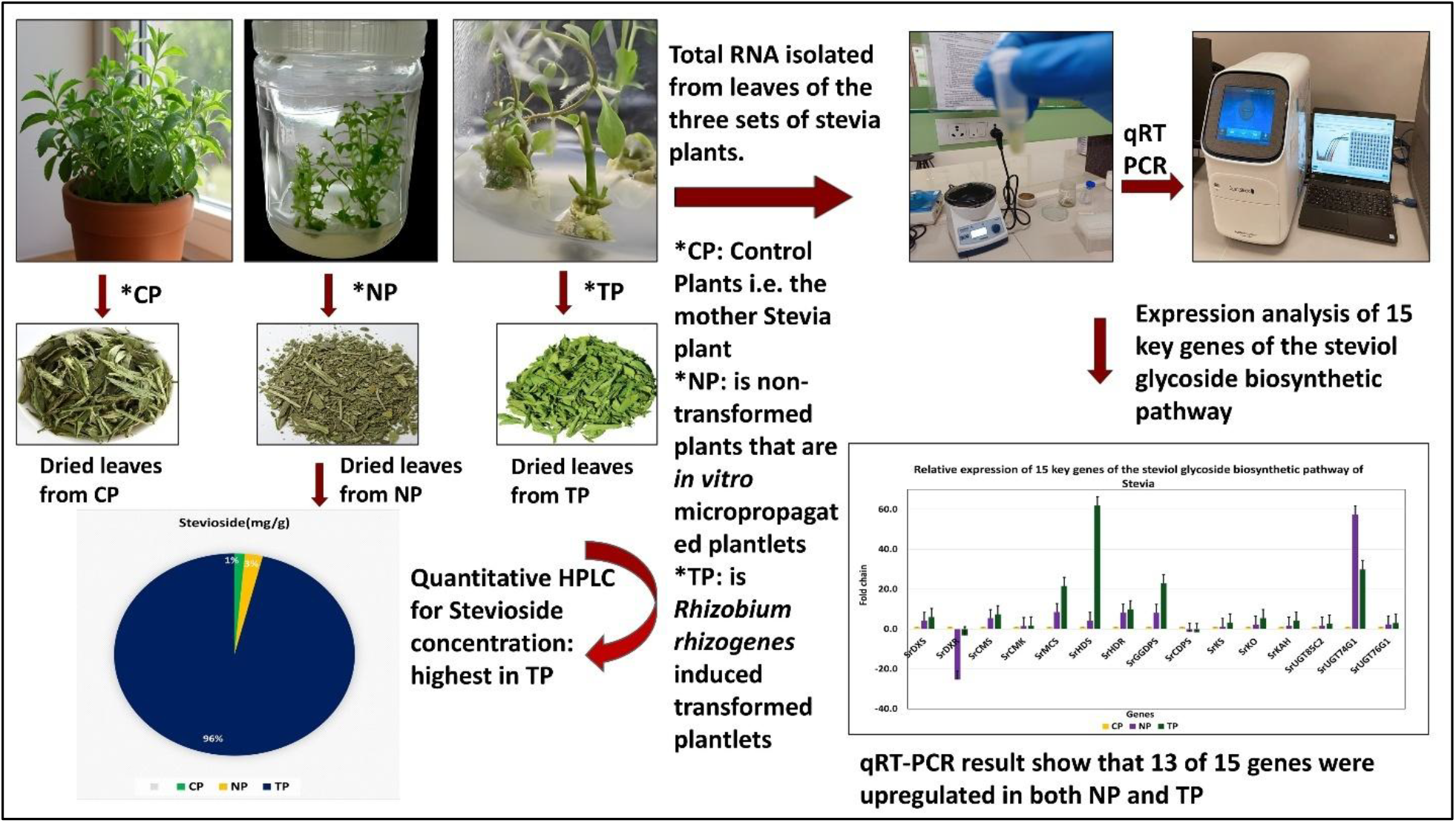

## 1. INTRODUCTION

*Stevia rebaudiana* (Bertoni), is a perennial shrub that belongs to the Asteraceae family of flowering plants. It is native to Paraguay and is noted for having a sweet taste due to a particular type of tetracyclic diterpenoids, called Steviol glycosides (SGs) found in the leaves [1]. These diterpenoids are naturally occurring, low-calorie sweeteners found in three species of plants, *Angelica keiskei*, Chinese *Rubus sauvissimus*, and *Stevia rebaudiana* [2]. SGs are primarily located in the leaves of Stevia plants and they are later transported to other parts of the plant [3]. To date, more than 60 SGs have been found in Stevia [4], [5]. The first glycoside to be extracted from Stevia leaves was Stevioside, which was then followed by the isolation of *rebA*, Dulcoside A, *rebB*, C, D, E, F, I, M, and Steviol-bioside [6]. *Stevia rebaudiana* typically contains Stevioside (5–10%), Rebaudioside A (2–5%), Rebaudioside C (1%), Dulcoside A (0.5%), and trace amounts of Rebaudiosides D, E, F (0.2%) and Steviol-bioside (0.1%) [7]. Among these, Stevioside and Rebaudioside A are the crucial SGs because they are the final products of the pathways. In addition to these secondary metabolites, plants also contain flavonoids, phenolics, and alkaloids that may have therapeutic benefits [8]. Stevia has many therapeutic properties such as, antidiabetic [9] antihypertensive, anticancer and antitumour effect [10] and antimicrobial effect [11].

Stevia is typically propagated using both seeds and shoot cuttings; however, seed propagation is less popular due to low production and germination efficiency, while shoot cutting propagation is a labour-intensive and time-consuming technique with a low success rate [12]. Optimum *in vitro* conditions are necessary for the development of an efficient method for mass propagation of Stevia. Thus, an alternative method such as plant tissue culture, that has high efficacy, is used to raise Stevia plants to elevate the quality and quantity of Steviol glycosides and further enhance the taste of *S. rebaudiana* [13],[14].

In the present study, plantlets generated through two distinct tissue culture approaches, micropropagation and *Rhizobium rhizogenes*-mediated transformation were utilized as the primary experimental material [15]. The steviol glycoside (SG) biosynthesis pathway in *Stevia rebaudiana* synthesizes compounds responsible for the formation of the natural sweetness in the Stevia plants via a complex sequence of enzymatic reactions. It involves a series of enzymatic reactions mediated by approximately 15 key genes as mentioned in Figure 5. The pathway begins with geranylgeranyl diphosphate (GGPP), which is converted by copalyl diphosphate synthase (CPS) and kaurene synthase (KS) to ent-kaurene. This is subsequently oxidized by kaurene oxidase (KO) and kaurenoic acid hydroxylase (KAH) to produce steviol, the aglycone backbone of SGs. In the final steps, UDP-glycosyltransferases (UGTs) like UGT85C2, UGT74G1, and UGT76G1 catalyse sequential glycosylation reactions to generate stevioside and various rebaudiosides [3],[16]. The coordinated activity of these 15 genes shapes the production of SGs, creating compounds like stevioside and rebaudioside A, which are crucial to the sweetness profile of Stevia. Understanding and manipulating these genes in biotechnological applications can probably enhance the yield and composition of steviol glycosides, potentially creating sweeter and more stable stevia products for commercial use.

Recent studies have highlighted the roles of specific genes in the pathway. For example, UGT76G1 is critical for synthesizing rebaudioside A, a desirable sweetener due to its reduced bitterness compared to Stevioside [17]. Modulating UGT enzyme expression can enhance specific SGs, allowing selective breeding or genetic engineering to produce high-yield, consumer-preferred sweeteners [18].

In light of the previous discussion, the present study is aimed to analyse the relative expression profiles of fifteen key genes involved in the steviol glycoside (SG) biosynthetic pathway in *Stevia rebaudiana* leaf tissue. Quantitative real-time PCR (qRT-PCR) was employed to assess relative gene expression across two plant groups: transformed plants (TP), non-transformed plants (NP), in comparison with *in vivo* control plants (CP) with the objective of developing a *Stevia* system exhibiting enhanced sweetness. To correlate gene expression with SG accumulation, quantitative analysis of steviol glycosides using stevioside as the reference standard was conducted via high-performance liquid chromatography (HPLC). This allowed for the comparative estimation of stevioside content among TP, NP, and CP plants, thereby validating the expression data obtained through qRT-PCR.

## 2. MATERIALS AND METHODS

### 2.1. Plant material

*Stevia rebaudiana* Bertoni (Family: *Asteraceae*) plants were procured from Sunrise Agro Nursery for Medicinal Plants (18°35’20”N, 73°46’30”E), Wakad, Pune. These *in vivo* plants were subsequently maintained and cultivated under controlled conditions in the greenhouse facility of Dr. D.Y. Patil Biotechnology and Bioinformatics Institute, Tathawade, Pune, and served as control group (CP) in the present study. The same plants also served as the source of explants for regeneration studies.

Specifically, internodal segments excised from CP plants were used for direct regeneration of non-transformed plantlets (NP) through micropropagation. For the generation of transformed plantlets (TP), micro-shoots derived from NP lines were co-cultivated with *Rhizobium rhizogenes* to induce hairy root formation. Hairy-root-bearing micro-shoots were then cultured to regenerate TP plantlets.

The regeneration protocols for both NP and TP plantlets were optimized at the Plant Tissue Culture Laboratory of Dr. D.Y. Patil Biotechnology and Bioinformatics Institute, Pune, and Rise N’ Shine Biotech Pvt. Ltd., Theur, Pune. The molecular confirmation of transformation through PCR amplification of *rolB, rolC*, and *virD2* genes was also performed which confirmed stable integration of *Ri* T-DNA into Stevia hairy roots and the corresponding regenerated plantlets [15]. These regenerated plantlets constituted the primary experimental material for the current study (**Figure 1**).

**Figure 1.**
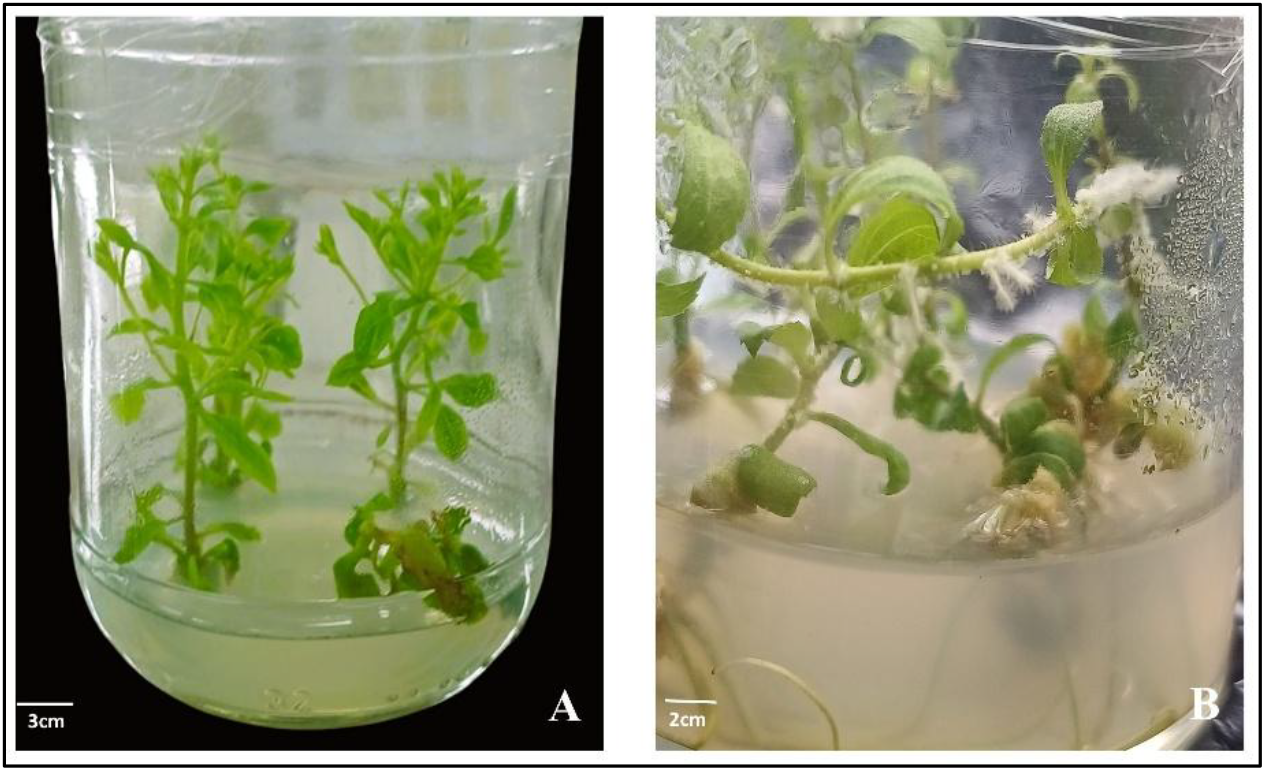
*Stevia rebaudiana* plant material. A) Micropropagated plantlets; B) *Rhizobium rhizogenes* mediated transformed plantlets [15].

### 2.2. Total RNA extraction and first-strand cDNA synthesis

Total RNA extraction was carried out in three replications from the stevia leaf tissue of CP, NP and TP using the Spectrum Plant Total RNA kit (SIGMA Life Sciences) by following the manufacturer’s protocol.

The extracted total RNA pooled (three replications together) of each treatment that were quantified using Nano-Drop spectrophotometer (Cytation5 Multimode Microplate Reader, BioTek) and run on an agarose gel (1%, w/v) electrophoresis. The cDNA was synthesized using the pooled RNA (3 replicates / treatments) sample of treatments using Go-Script Reverse Transcription System (Promega Biotech India Pvt Ltd) by following the manufacturer’s protocol.

### 2.3. Selection of Steviol Glycoside pathway genes and primer designing

A total of fifteen key genes involved in the steviol glycoside biosynthesis pathway were selected based on their known roles in the metabolic conversion of precursors into major steviol glycosides, primarily stevioside and rebaudioside A [4]. The nucleotide sequences of all these genes were retrieved from the NCBI GenBank database (https://www.ncbi.nlm.nih.gov). The selection was based on sequence availability, functional annotation, and previous reports [4] on their involvement in SG biosynthesis in *Stevia rebaudiana*. Corresponding accession numbers for each gene are provided in **Table 1**. Primer pairs were designed using Primer-BLAST (NCBI) and Primer3Plus (https://www.primer3plus.com) under default parameters to ensure target specificity, optimal melting temperatures, and amplicon sizes suitable for qRT-PCR analysis that are mentioned in **Table 1**.

**Table 1.**
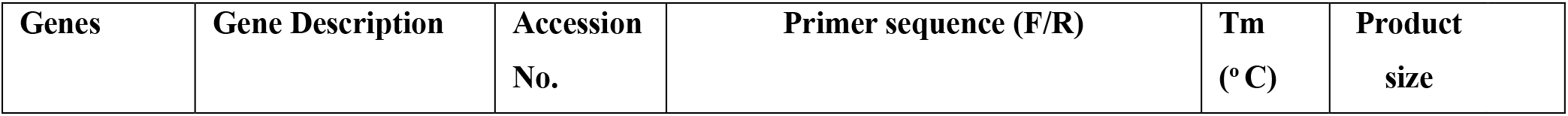
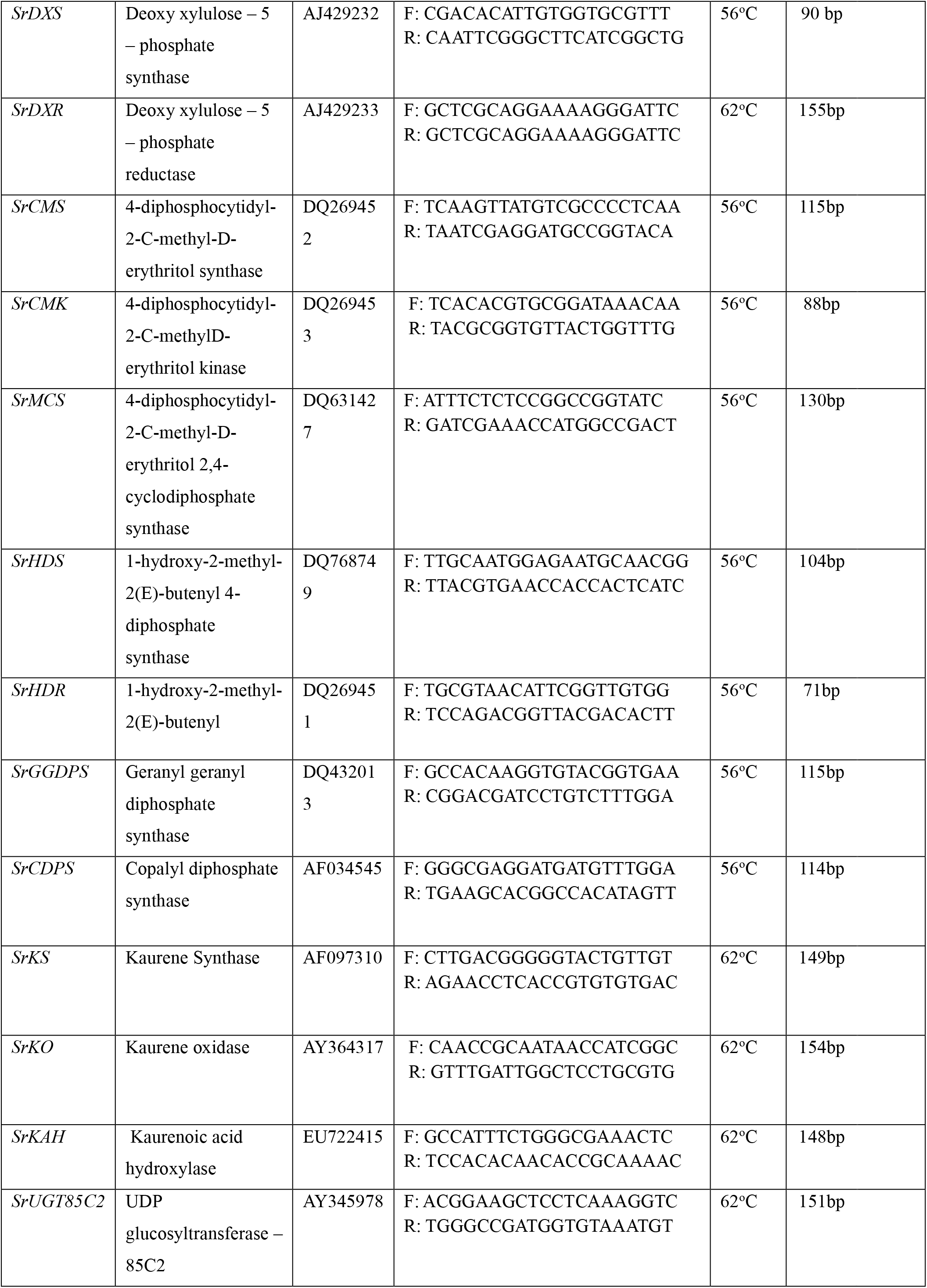
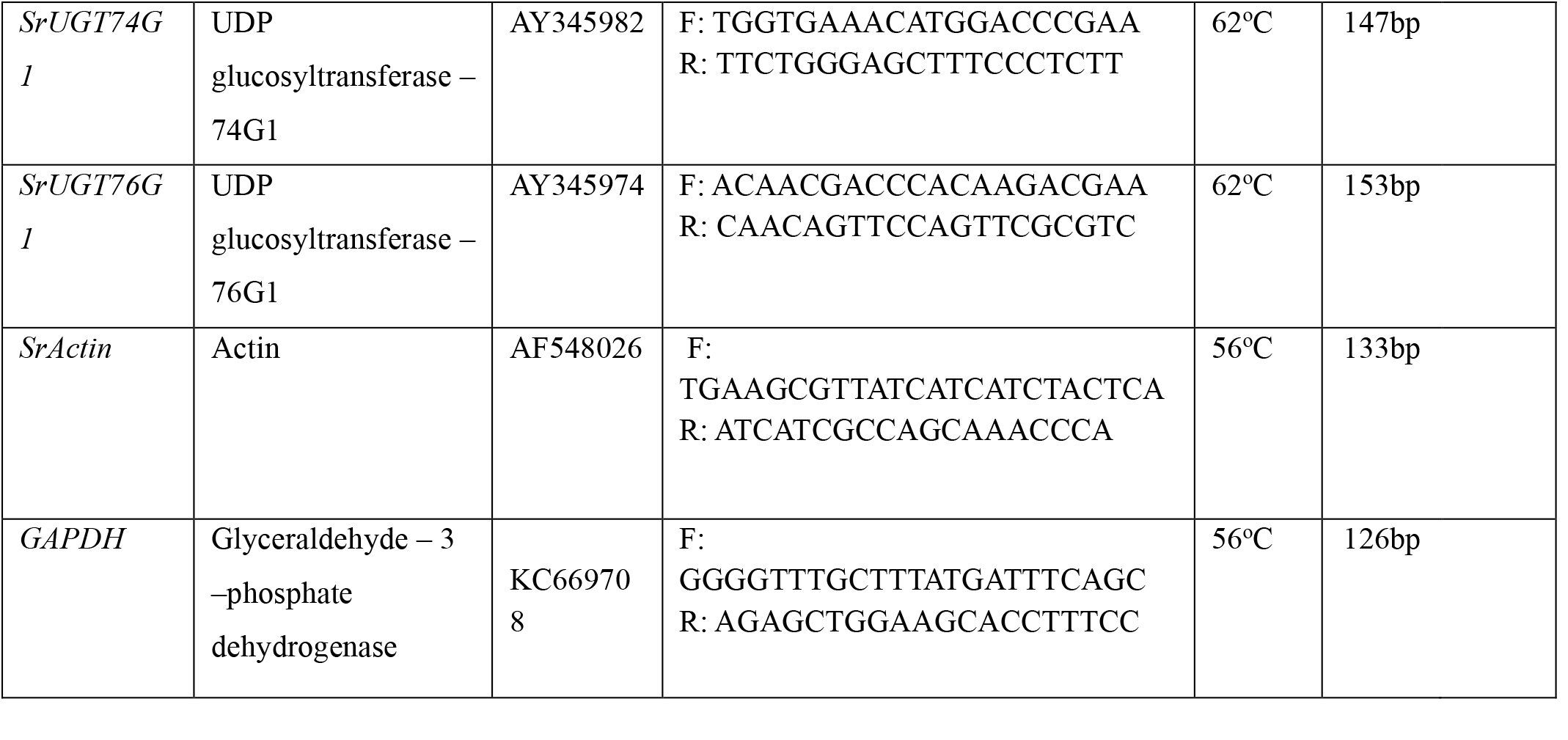
Primer sequences of 15 genes of Steviol glycosides (SGs) pathway and reference genes.

### 2.4. Gene expression profiling

The gene expression profiling of selected fifteen genes were performed using Quantitative real-time polymerase chain reaction (qRT-PCR). The two genes, *Actin* and *GAPDH* were used as a set of reference genes or internal control genes. The qRT-PCR reactions were carried out in Thermal Cycler1000 Touch CFX96 Touch™ (Bio-Rad) using iTaq Universal SYBR green Supermix (Bio-Rad). Each of the reactions was performed in triplicate containing SYBR green Supermix (5ul), template cDNA (1 μl), each of the primers (0.4μl, 10μM), and RNAse-free water (3.2 μl) with 10 μl total volume.

The qRT-PCR profile for two reference genes and eight Steviol Glycoside pathway genes (includes *CMS, CMK, MCS, HDR, GGDPS, CDPS, HDS* and *DXS*) were as follows, 1min (95°C), 10s (95°C) for 40 cycles, 30s (56°C), 5s (65°C), 5s (95°C) and for other nine genes (*KO, KS, KAH, DXR, UGT76G1, UGT74G1* and *UGT85C2*) as follows, 1min (95°C), 10s (95°C) for 40 cycles, 30s (62°C), 5s (65°C), 5s (95°C) for fluorescent signal recording followed by high resolution melting curve obtained after the cycle 95°C (15s), with constant increment in the temperature from 65°C (15 s) and 95°C (1s). Melting curve analysis was done and internal control genes *SrActin* and *GAPDH* were used to normalise the data individually. Finally, data was analysed using the software, Bio-Rad CFX maestro version 2.3.

The fold change values from the gene expression profiling of genes in leaf tissue of CP, NP and TP were calculated using 2-^ΔΔCT^ method [19] and also heatmaps plot depicted in **Figure 5** using, Multi Experiment Viewer (MeV v4.9.0) software [20]. The fold change values of each gene from NP and TP were plotted in the graph as shown in **Figure2-4**.

Although CP (control plants), NP, and TP (*in vitro*-regenerated plantlets) were cultivated under distinct environmental conditions, leaf samples from both *in vivo* and *in vitro* plants were harvested after 1.5 months of growth to ensure comparable physiological status at the time of collection. The selection of *in vivo* plants as a reference for gene expression analysis aligns with prior transcriptomic studies conducted in rice and *Eucommia ulmoides* [21], [22]. Quantitative gene expression data were normalized using two stable internal reference genes, *SrActin* and *GAPDH*.

### 2.5. HPLC Analysis

Stevioside content in the leaf samples of CP, NP, and TP was quantified using High-Performance Liquid Chromatography (HPLC). Dried leaf powder was extracted with HPLC-grade methanol, defatted with hexane, and re-dissolved in acetonitrile. Filtrates were injected into an Agilent HPLC system equipped with a Zorbax Eclipse XDB–C18 column and detected at 204 nm. The mobile phase consisted of methanol and 0.05% o-phosphoric acid (75:25, v/v) at pH 3.15. Chromatographic data were analysed using EZChrom Elite software. Stevioside concentrations were calculated against a standard curve and expressed in mg/g dry weight.

### 2.6. Statistical analysis

The Quantification studies were run in six replications for each leaf sample from CP, NP and TP. The result was expressed as the mean value of Stevioside concentration ± standard error. SPSS (Statistical Package for Social Science; version 17) was used for the statistical analysis. To assess the significance of the mean values at (p<0.05) the one-way ANOVA test was carried out and for pairwise comparison, the Post Hoc test (Tukey’s HSD) was applied.

## 3. RESULTS

### 3.1. Quantitative real time PCR (qRT-PCR) analysis

In the present study, the expression profiling (qRT-PCR) of key genes (fifteen genes) involved in the Steviol Glycoside biosynthetic pathway were studied from leaf tissues of *in vitro* regenerated non-transformed plants (NP), *in vitro* transformed plants (TP) and control plants (CP) grown under environmental conditions.

The expression of *SrDXS* (Deoxyxyulose-5-phosphate synthase) exhibited higher in *in vitro* transformed plants (TP) (5.0-fold change, up-regulated) followed by *in vitro* regenerated non-transformed plants (NP) (3.0-fold change, up-regulated) as compared to the control plant (CP) as shown in **Figure 2(a)**. The *SrDXR* (deoxyxyulose-5-phosphate reductase) gene exhibited lower expression in TP (−3.2-fold change, down-regulated) and in NP (−25.4-fold change, highly down-regulated) compared with CP **[Figure 2(a)]**.

**Figure 2.**
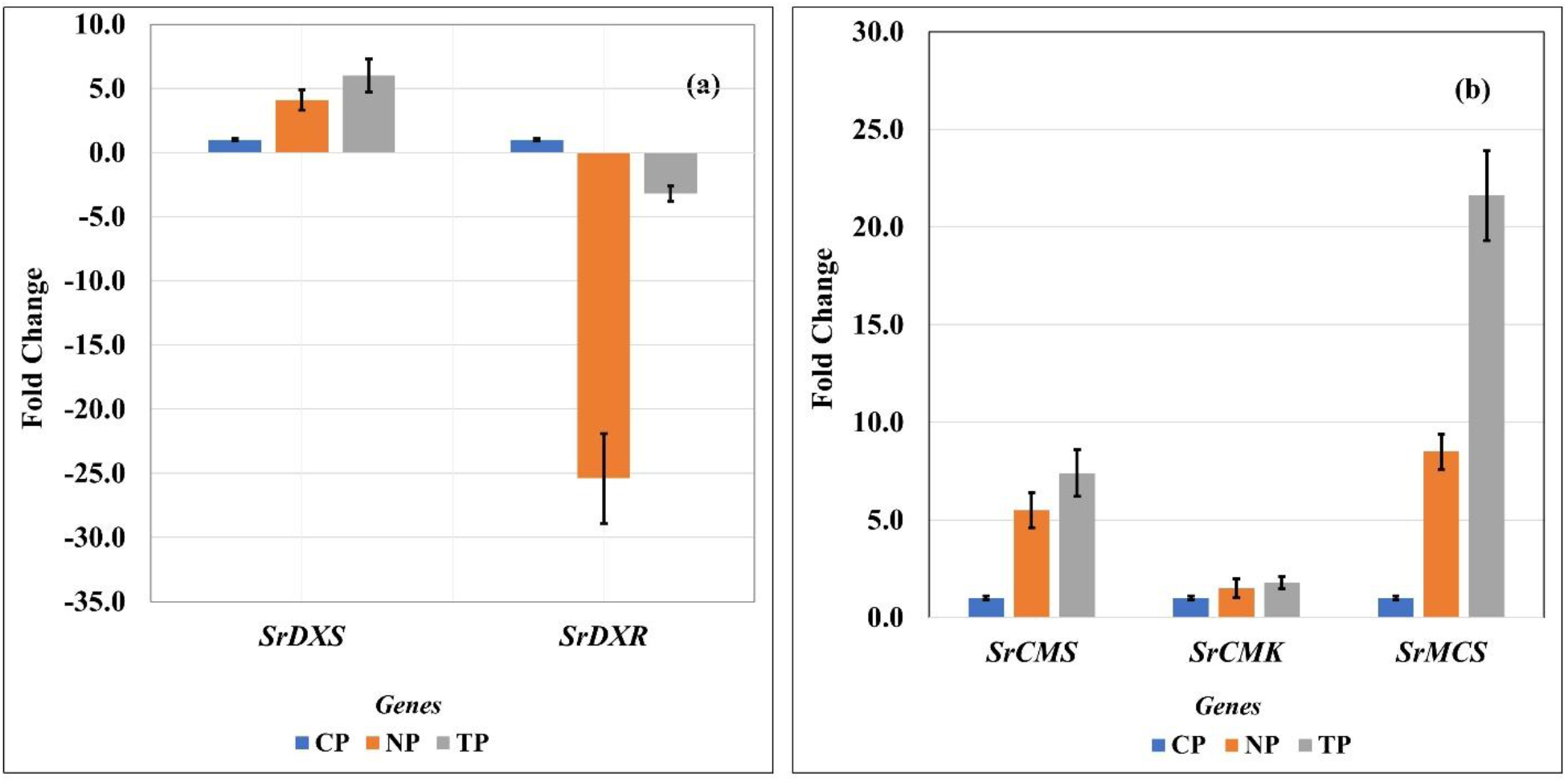
Gene expression profiling of Steviol Glycoside biosynthesizing genes **(a)** *SrDXS* and *SrDXR*, **(b)** *SrCMS, SrCMK* and *SrMCS* in leaves tissues of TP, NP and CP. Error bars on the top indicate standard deviation of three technical replicates. *SrActin* and *GAPDH* served as two reference genes.

The transcript abundance of three diphosphocytidyl related genes, *SrCMS* (4-Diphosphocytidyl-2-C-methyl-d-erythritol synthase), *SrCMK* (4-Diphosphocytidyl-2-C-methyl-d-erythritol kinase) and *SrMCS* (4-Diphosphocytidyl-2-C-methyl-d-erythritol 2,4-cyclodiphosphate synthase) exhibited higher expression in TP (7.4, 1.8 and 21.6-fold change respectively, up-regulated) followed by NP (5.5, 1.5 and 8.5-fold change respectively, up-regulated) when compared with CP as shown in **Figure 2(b)**.

Furthermore, the transcript level of two 1-Hydroxy-2-methyl-2(E)-butenyl-4-diphosphate synthase/reductase genes, *SrHDS* (1-Hydroxy-2-methyl-2(E)-butenyl-4-diphosphate synthase) *SrHDR* (1-Hydroxy-2-methyl-2(E)-butenyl-4-diphosphate reductase) exhibited the highest increase in TP (61.9

-fold change, highly up-regulated and 9.9-fold change respectively, up-regulated) and in NP (4.1 and 8.2-fold change respectively, up-regulated) as compared to CP as shown in **Figure 3(a)**.

**Figure 3.**
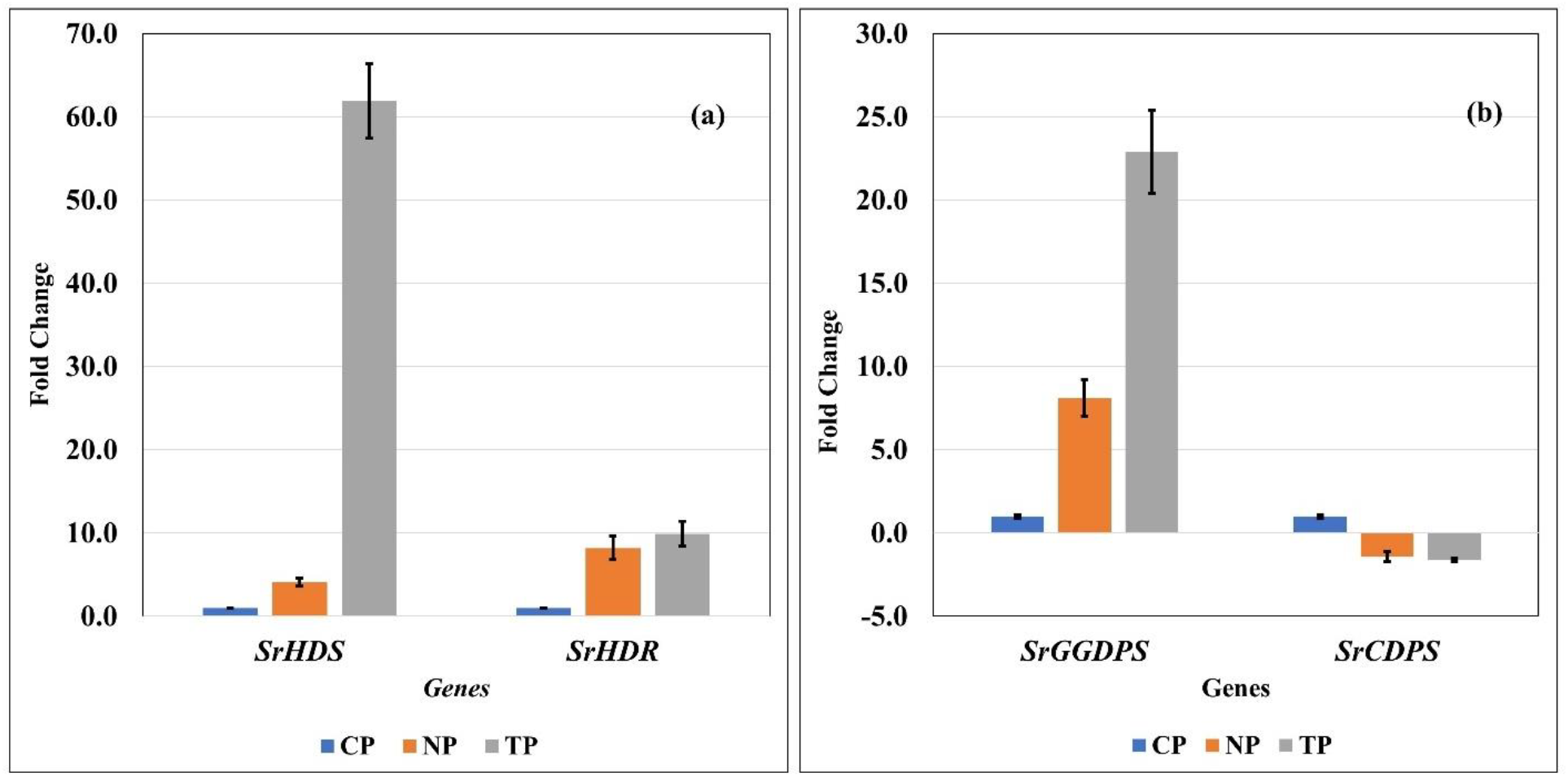
Gene expression profiling of Steviol Glycoside biosynthesizing genes **(a)** *SrHDS* and *SrHDR*, **(b)** *SrGGDPS* and *SrCDPS* in leaves tissues of TP, NP and CP. Error bars on the top indicate standard deviation of three technical replicates. *SrActin* and *GAPDH* served as two reference genes.

Among the two diphosphate synthase genes, *SrGGDPS* (Geranylgeranyl diphosphate synthase) gene showed the highest expression in TP (22.9 -fold change, highly up-regulated) followed by NP (8.1-fold change, up-regulated) as compared to CP, shown in **Figure 3(b)**. Whereas in contrast, *SrCDPS* (Copalyl diphosphate synthase) gene exhibited low expression in TP (−1.6-fold difference, that is down-regulated), followed by NP (1.4-fold change, down-regulated) in comparison with CP **[Figure 3(b)]**.

Similarly, the transcript level of two genes, *SrKS (*Kaurene synthase) and *SrKO (*Kaurene oxidase) exhibited higher in TP (3.3 and 1.1-fold increase respectively, up-regulated) followed by NP (5.5 and 2.2 -fold change respectively, up-regulated). However, the higher expression of *SrKAH* (Kaurenoic acid hydroxylase) was recorded in TP (4.2-fold change, up-regulated) and NP (1.7-fold change, up-regulated) as shown in **Figure 4(a)**.

**Figure 4.**
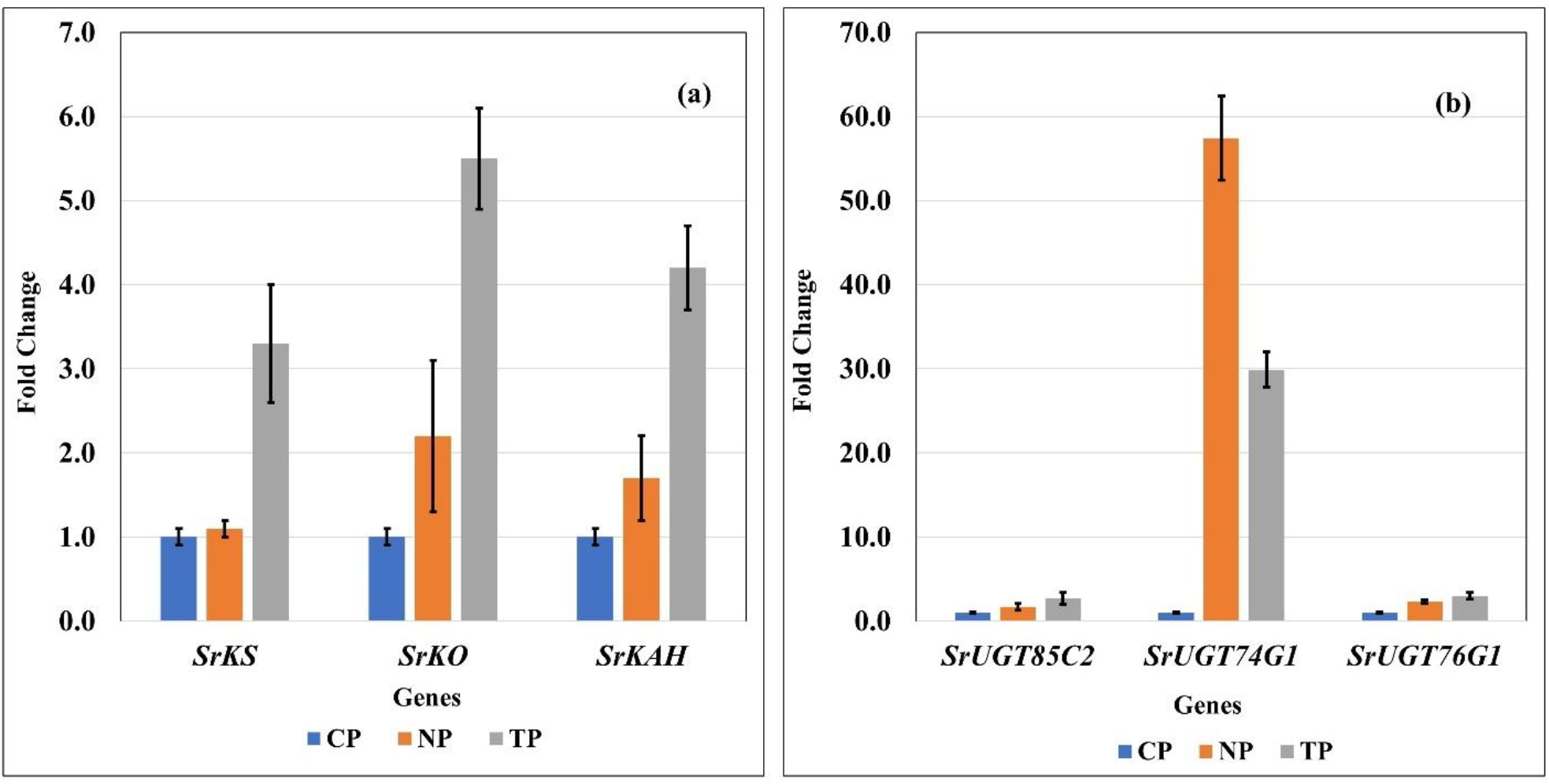
Gene expression profiling of Steviol Glycoside biosynthesizing genes **(a)** *SrKS* and *SrKD* and *SrKAH*, **(b)** *SrUGT85C2, SrUGT74G1* and *SrUGT76G1* in leaves tissues of TP, NP and CP. Error bars on the top indicate the standard deviation of three technical replicates. *SrActin* and *GAPDH* served as two reference genes.

**Figure 5.**
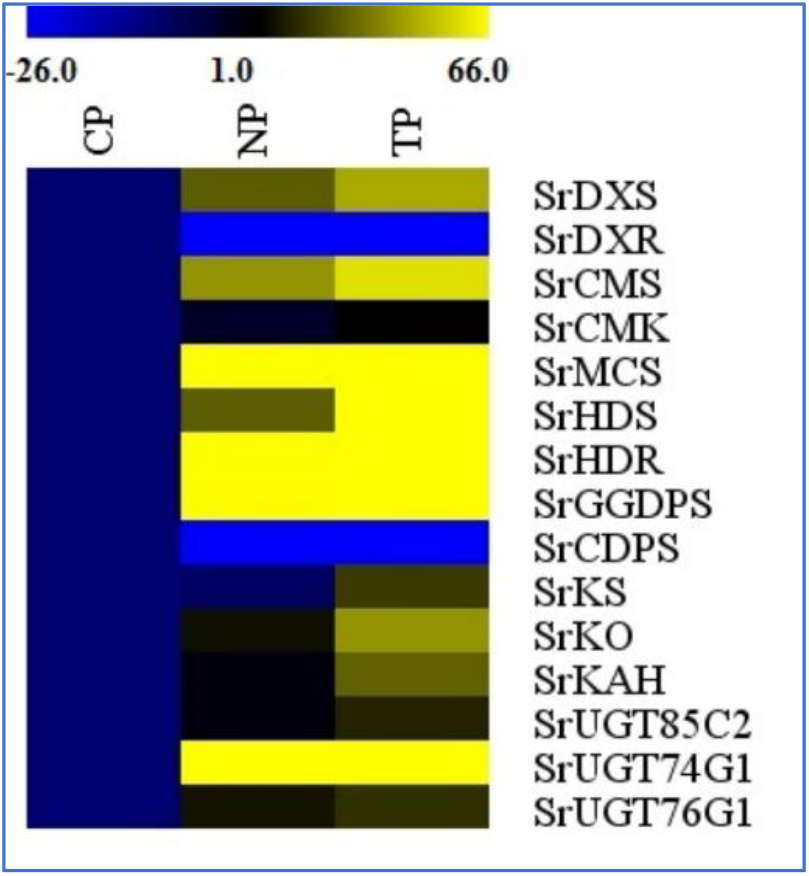
Heatmap representation of fifteen steviol glycoside biosynthesis pathway genes expression profile in leaf tissue of TP, NP and CP. The colour scale at the top show expression level (fold change) values. Yellow indicates a higher expression level (up-regulation), and blue indicates a lower expression level (down-regulation) of genes. For, CP the colour remains uniform throughout.

Furthermore, among the glucosyltransferase genes, *SrUGT85C2 (*UDP glucosyltransferase-85C2) and *SrUGT76G1 (*UDP glucosyltransferase-76G) were highly expressed in TP (2.7 and 3.0 -fold difference respectively, up-regulated) followed by NP (1.7 and 2.3 -fold change, up-regulated) when compared with CP, shown in **Figure 4(b)**. However, *SrUGT74G1* (UDP glucosyltransferase-74G) gene exhibited higher expression in *in vitro* regenerated non-transformed plants (NP) about 57.4-fold change (highly up-regulated), followed by *in vitro* regenerated transformed plants (TP), about 29.9-fold change (highly up-regulated) in comparison with CP **[Figure 4(b)]**.

Thus, comparatively higher expression profiling of SGs genes in *in vitro* transformed plants i.e. TP (plantlets regenerated from induced hairy roots *via. Rhizobium rhizogenes* transformation) indicates that the modulation occurred in genes involved in the Steviol Glycoside biosynthetic pathway might be leads to enhance the production and content of Steviol glycosides.

### 3.2 HPLC analysis

The key feature of *Stevia* is its steviol glycoside accumulation. Examination of Stevioside concentrations of leaf tissues of CP, NP and TP is carried out by HPLC. According to the results shown in **Table 2**, the highest amount of stevioside was produced by *Rhizobium rhizogenes*-mediated transformed plantlets. It can be concluded that transformation increases steviol glycoside contents in the leaves of stevia. One-way ANOVA shows that all the data are significant with *p* value < 0.05. Tukey’s HSD test, showed that all pairwise comparisons (CP vs NP, CP vs TP, NP vs TP) are statistically significant (p < 0.05). This confirms that stevioside concentration significantly differs among CP, NP, and TP, with TP showing the highest stevioside concentration. The corresponding HPLC chromatograms illustrating peak profiles of stevioside from CP, NP, and TP leaf extracts are presented in **Figure 6**.

**Table 2.** Stevioside concentration in mg/g of the leaf extracts of the *Stevia rebaudiana* Control plants CP, non-transformed plants NP and transformed plants TP.

| Sample | Retention time | Area | Area % | Height | Stevioside(mg/g) [Mean ± SE] |
| --- | --- | --- | --- | --- | --- |
| CP | 6.273 | 193133 | 100.00 | 23026 | 0.024± 0.02 <sup>a</sup> |
| NP | 6.217 | 317420 | 100.00 | 39609 | 0.042± 0.01 <sup>b</sup> |
| TP | 6.861 | 304175 | 100.00 | 25401 | 1.67± 0.13 <sup>c</sup> |

\* Values are the mean of Stevioside concentration in different set of plants determined by one-way ANOVA that shows data are statistically significant ( $p < 0.05$ ). Post Hoc test of pairwise comparison was done via Tukey HSD. Different letters represent statistically significant data. The experiment was performed in 6 replicates.

**Figure 6.**
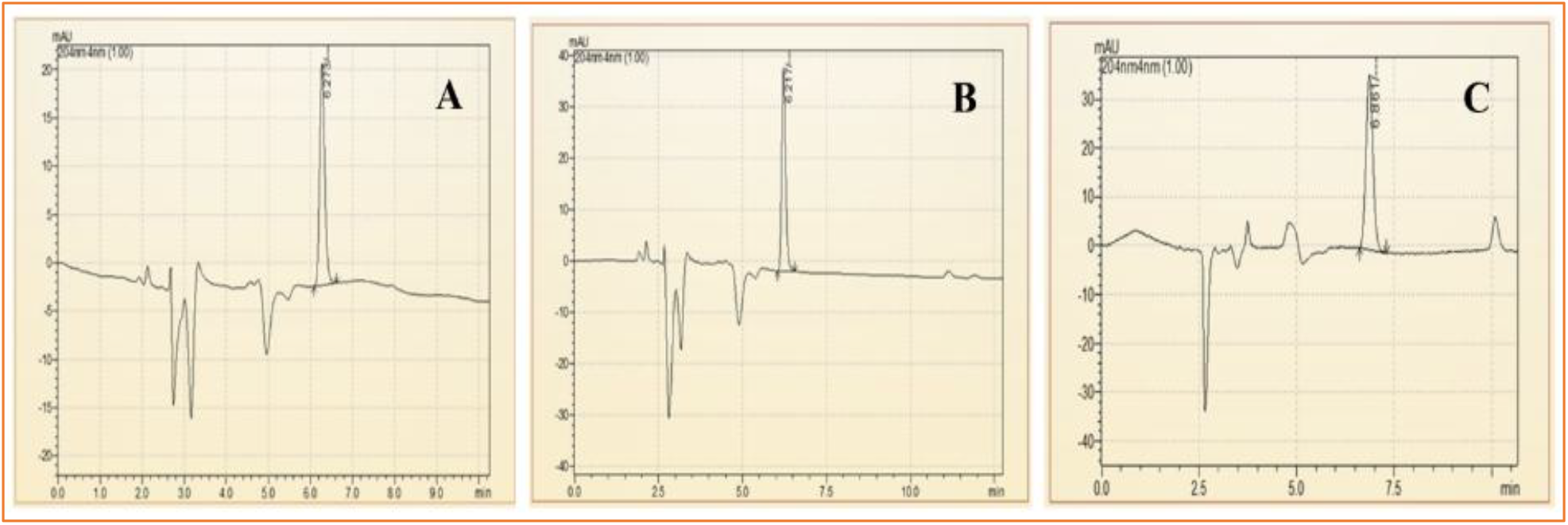
HPLC chromatogram of the leaf extracts of the Stevia Control plants CP, non-transformed plantlets NP and transformed plantlets TP.

## 4. DISCUSSION

In the current study, two types of transcript accumulation patterns, up-regulation and subsequently varying rates of down-regulation, were seen in the genes of the steviol glycoside biosynthesis pathway in *Stevia rebaudiana* Bertoni, both in transformed and non-transformed lines. The highest transcript levels among the up-regulated genes were found in the leaves of the transformed plants, indicating that, out of the 15 pathway genes investigated in this study, *Rhizobium rhizogenes*-infected plants had superior transcript level, in general. The present findings are quite similar to the findings of Sarmiento-López, 2020; Libik-Konieczny *et al*, 2020 and Sanchéz-Cordova *et al*, 2019 [23], [24], [25]. However, specifically 13 genes were upregulated and two of them were downregulated in the current study.

In our findings, higher transcript abundance (i.e. up-regulation) in three genes, *SrKS* (Kaurene synthase), *SrKO* (Kaurene oxidase) and *SrKAH* (Kaurenoic acid hydroxylase) were recorded in TP and NP as compared to CP, which showed enhanced SGs content in leaves corroborated with the earlier finding in Stevia by Zheng *et al*, 2019 and Nasrullah *et al*, 2023 [26], [27]. The genes *SrKO* and *SrUGT74G1*show high expression, leading to an increase in Stevioside level. This finding correlates with studies by Nasrullah *et al*, 2023 in *Stevia rebaudiana*, demonstrating their significant role in enhancing SG content [27].

In the present research, all NP and TP lines were propagated using the same combination of plant hormones in MS media and maintained under the same growth conditions [15], however, a higher *SrUGT74G1* expression level was noted in non-transformed plantlets (NP) when compared with transformed lines (TP). Expression increase of *SrUGT74G1* in NP could be due to metabolic reprogramming during micropropagation stress. The lack of comparable expression in TP, even with the same culture conditions, may indicate that in secondary metabolism of plants, the feedback regulation mechanisms responsive to the complete metabolites tend to inhibit the expression of biosynthetic pathways. For instance, Gachon *et al*, 2005 and Tiwari *et al*, 2016 discuss how glycosylation, mediated by glycosyltransferases plays a crucial role in regulating hormone homeostasis and secondary metabolite biosynthesis. [28],[29].

In TP lines, higher flux through the steviol biosynthetic pathway might activate such feedback loops, leading to the downregulation of *SrUGT74G1*. Conversely, NP plants, with comparatively lower precursor flux, may not trigger these feedback mechanisms, resulting in higher expression levels of *SrUGT74G*. Another reason might be the choice of explants [30]. For the NP plants, nodal explants were used for direct shoot regeneration while, in case of TP plants, micro-shoots with hairy roots were used as explants for transformed plantlet regeneration [15]. Additionally, post-transcriptional regulation, including mechanisms mediated by microRNAs (miRNAs), can influence gene expression levels. Kajla *et al*, 2023 highlight the role of miRNAs in fine-tuning the expression of genes involved in secondary metabolite biosynthesis. Such regulatory processes can lead to variations in gene expression independent of genetic transformation [31].

In the present study, two genes, *SrDXS* and *SrCDPS* indicated a downregulation and despite that, in *in-vitro* generated plants, stevioside levels were still high, as confirmed by HPLC analysis. This could be achievable due to such genes being part of complex regulatory network in the SG pathway, which may involve a feedback mechanism, post-transcriptional regulation, and compensatory upregulation of other genes to sustain or even elevate overall steviol glycoside production. The increased flux through the SG pathway in *in-vitro* plantlets may trigger feedback regulation mechanisms. These mechanisms, which respond to the total metabolites, prefer to suppress the expression of biosynthetic pathways. On the other hand, *in vivo* plants, with relatively reduced precursor flux, might not activate these feedback mechanisms, resulting in varying expression levels. Moreover, the upregulation of major downstream UDP-glycosyltransferase (UGT) genes, specifically *SrUGT76G1*, could counteract the downregulation of upstream genes such as *SrCDPS* and *SrDXR*, resulting in net increase in steviol glycoside accumulation. Similar trends have been reported in Stevia transformation studies, where enhanced glycosylation contributed to higher SG yields despite variations in early pathway gene expression [32], [33]. Lower expression of these genes can also act as positive regulators of the pathway. The correlation between gene expression and SG quantification highlights the complex regulation of the biosynthetic pathway. While some genes exhibit downregulation, the overall metabolic flux toward stevioside biosynthesis appears to be driven by the enhanced expression of key glycosyltransferases. This underscores the potential for targeted genetic modifications to optimize SG production in Stevia [34], [35].

HPLC data of the current research showed that transformed *Stevia* plants had higher stevioside content compared to micropropagated plants (NP), which in turn accumulated more stevioside than the *in vivo* -grown control plants (CP). This trend largely coincides with the gene expression data obtained through Real-Time PCR analysis in the current study. Similar findings regarding enhanced SG accumulation in *Rhizobium rhizogenes*-mediated transformed *Stevia rebaudiana* plantlets were reported by Sánchez-Córdova and co-workers in (2019) [25]. The rise in stevioside content in NP as compared to the control plants might be attributed to tissue culture-induced metabolic reprogramming [36], where the controlled *in vitro* environment and synchronized developmental stage of regenerated plantlets can enhance secondary metabolite biosynthesis.

The present work mainly focused on HPLC analysis to quantify stevioside, one of the most predominant and important steviol glycosides found in *Stevia rebaudiana*. Stevioside is a central intermediate in the biosynthetic pathway and is frequently used as a reliable metabolic marker owing to its high accumulation in the leaf tissues. If other glycosides such as rebaudioside A were to be measured, a better picture of the metabolite spectrum would have emerged; however, the targeted analysis of stevioside alone provides a strong and representative estimate of the biosynthetic activity. This holds true for the main objective of correlating gene expression patterns with the core output of the steviol glycoside biosynthesis pathway. Additionally, focusing on stevioside has allowed for an accurate assessment of the impact of transformation and regeneration conditions on its accumulation.

## 5. CONCLUSION

In the present study, thirteen out of the fifteen analysed genes involved in the steviol glycoside (SG) biosynthesis pathway exhibited higher expression in the leaves of in vitro transformed Stevia plants (TP) compared to in vitro regenerated non-transformed plants (NP) and control plants (CP). This upregulation suggests a positive correlation between gene expression and enhanced biosynthesis and accumulation of steviol glycosides in leaves. Furthermore, HPLC analysis confirmed that transformed plantlets demonstrated a higher stevioside content, indicating their potential reliability for improved steviol glycoside production. These transformed plant lines could serve as an alternative approach for large-scale production of steviol glycosides, facilitating their evaluation in animal trials for prospective commercial applications as a natural sweetener. Steviol glycosides, being a plant-derived, low-calorie sugar substitute, could be particularly advantageous for individuals managing metabolic disorders, including high blood sugar, cardiovascular issues, and excessive weight gain. Their widespread application in the food industry may contribute to the development of healthier dietary alternatives, promoting overall well-being. This study highlights the potential for molecular-level manipulation of the SG biosynthetic pathway to enhance steviol glycoside production. However, further molecular studies and pathway exploitation are essential to fully harness the biotechnological potential of Stevia rebaudiana for commercial applications.

## 6. ACKNOWLEDGEMENTS

Authors are grateful to Dr. D. Y. Patil Biotechnology and Bioinformatics Institute, Dr, D, Y, Patil Vidyapeeth (Deemed to be University), Pune for providing the laboratory facilities and financial support. We are thankful to DST, NCL and Dr. D. Y. Patil pharmacy college, Pune for the instruments they provided for the current research.

## 7. AUTHOR CONTRIBUTIONS

All authors contributed significantly to the conception and design of the study, data acquisition, and/or analysis and interpretation of the results. Each author participated in drafting the manuscript and/or critically revising it for important intellectual content. All authors reviewed and approved the final version of the manuscript for submission to the journal and accept full responsibility for the content and integrity of the work.

## 8. AVAILABILITY OF DATA AND MATERIALS

All the data pertaining to this study, are in the possession of the authors and will be supplied upon request.

## 9. DECLARATIONS

### Compliance with ethical standards; Ethical issues

None.

### Consent for publication

All authors agree to be published

## 10. USING ARTIFICIAL INTELLIGENCE (AI) ASSISTANCE

The authors uphold that no artificial intelligence (AI) assisted technologies were used in the writing, editing, or preparation of this manuscript. Additionally, all images included in the manuscript are original and were not generated or altered using AI tools.

