## Supplemental Table 1 for "Steviol Glycoside biosynthesis pathway gene expression profiling of transformed and non-transformed plant leaf tissues of *Stevia rebaudiana* (Bertoni)": genes, curves and peak, Tm values.docx

Supplementary S1. Melt Curve Peak Temperatures (Tm) of the fifteen genes involved in steviol glycoside biosynthesis in *Stevia rebaudiana* and the two reference genes (*Sr Actin* and *Sr GAPDH*), obtained from qRT-PCR analysis.

The melt curve analysis presented in Supplementary S1 was conducted for three experimental groups:

- CP - Control Plants (*In vivo*)
- NP - Non-transformed Plantlets (regenerated via *in vitro* micropropagation)
- TP -Transformed Plantlets (regenerated from hairy root cultures using *Rhizobium rhizogenes* MTCC 532)

Each gene was analysed using three biological replicates per group (CP, NP, TP). The Tm values listed represent consistent amplification across all replicates, confirming the specificity and reproducibility of each primer set.

**1. *Sr DXS***

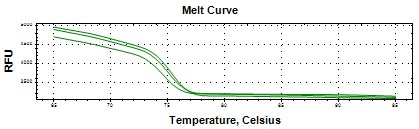

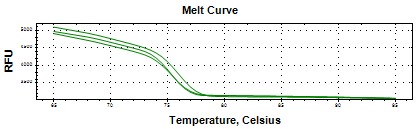

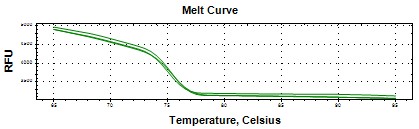

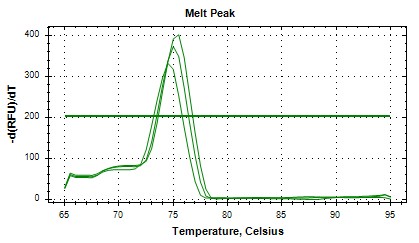

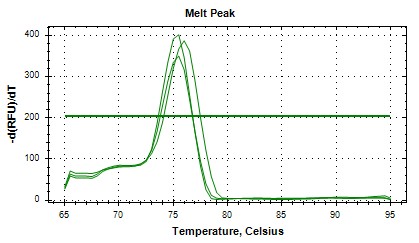

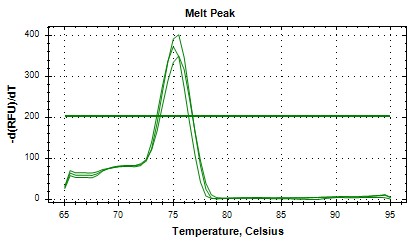

(CP) Tm =75	(NP) Tm =76.5	(TP) Tm =75.5

*Peak 1 likely main amplicon; others may reflect sequence variants or conformations

**2. *Sr DXR***

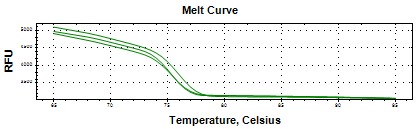

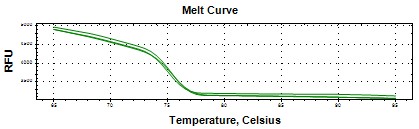

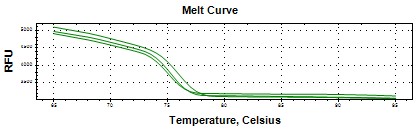

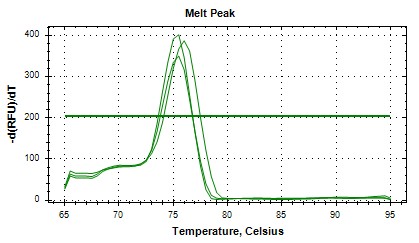

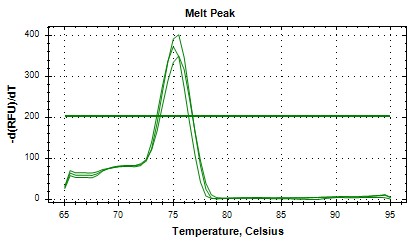

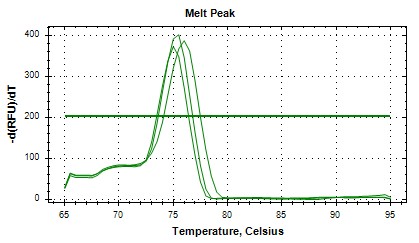

(CP) Tm =76.5	(NP) Tm =75.5	(TP) Tm =77

*Consistent peaks; indicates specific amplification

**3*. Sr CMS***

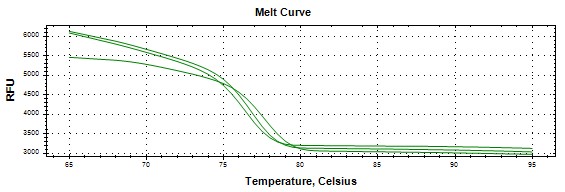

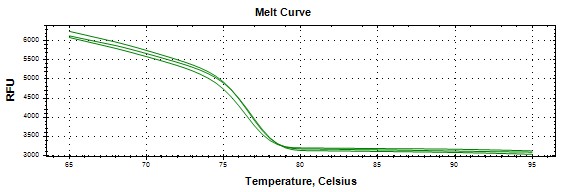

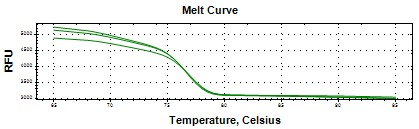

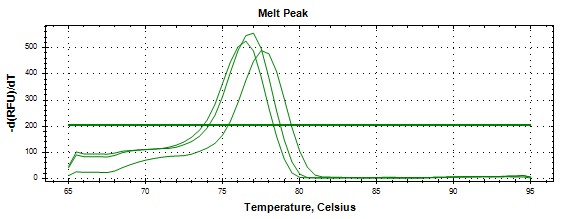

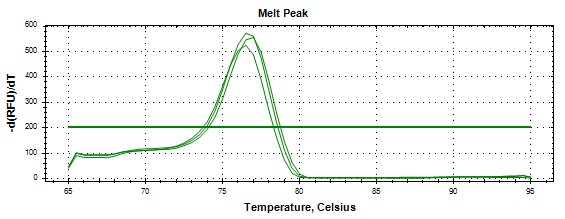

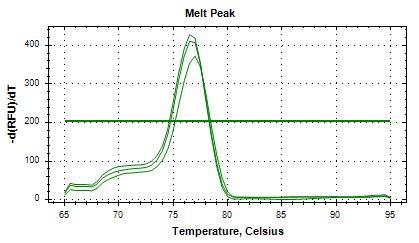

(CP) Tm =76.5	(NP) Tm =77	(TP) Tm =76

*All peaks within 1°C range, suggests specific target

**4*. Sr CMK***

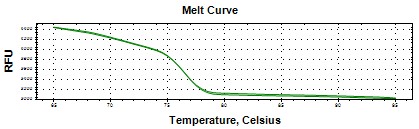

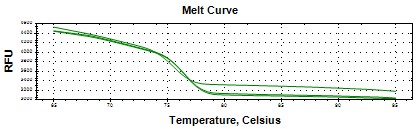

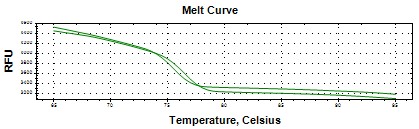

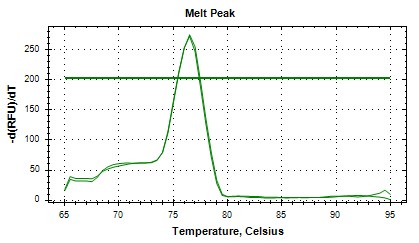

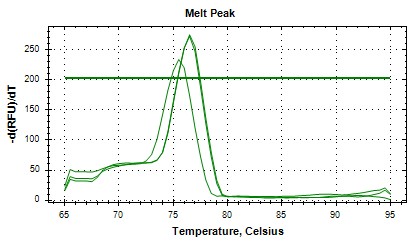

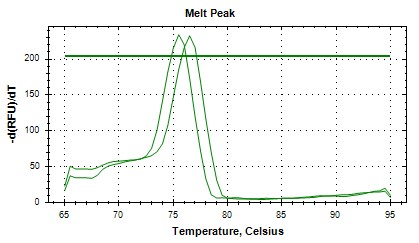

(CP) Tm =77	(NP) Tm =75.5	(TP) Tm =78

*Peak 3 slightly higher; may reflect GC-rich region

**5*. Sr MCS***

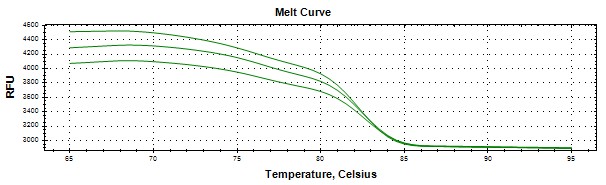

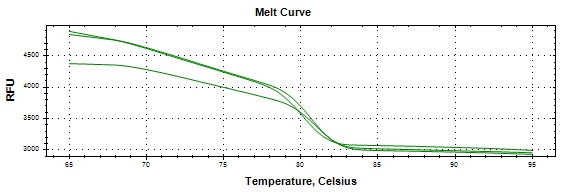

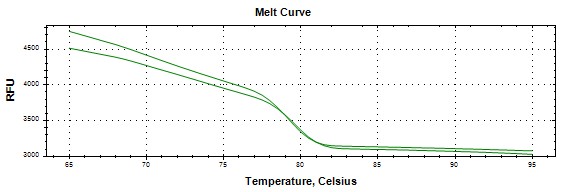

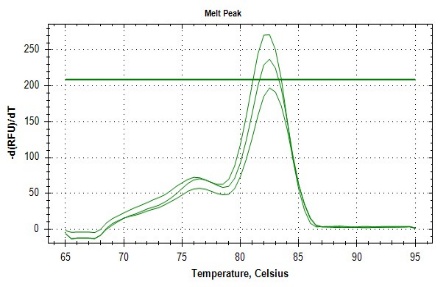

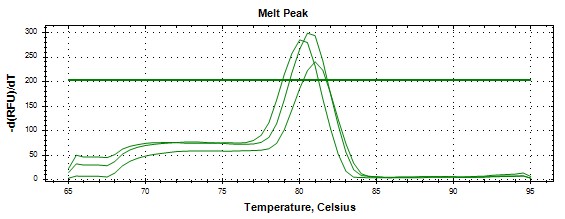

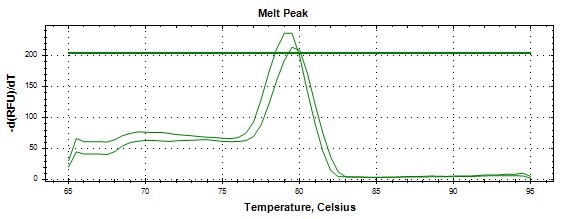

(CP) Tm =85	(NP) Tm =82.5	(TP) Tm =79.5

* Higher Tm; indicates GC-rich or longer product

**6. *Sr HDS***

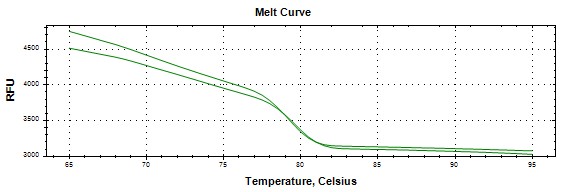

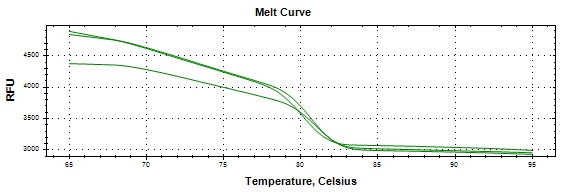

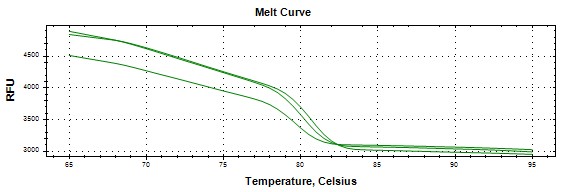

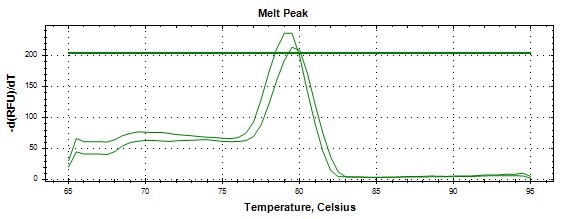

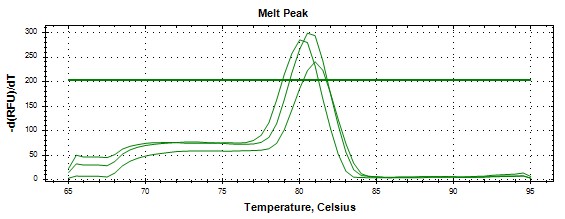

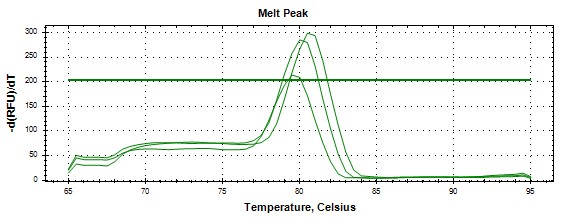

(CP) Tm =78.5	(NP) Tm =83.5	(TP) Tm =82

*Peak 2 dominant, likely correct product

**7*. Sr HDR***

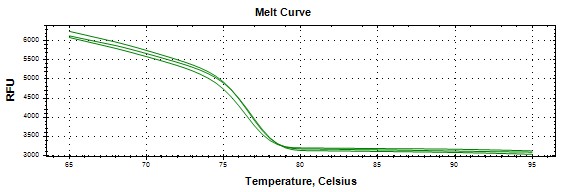

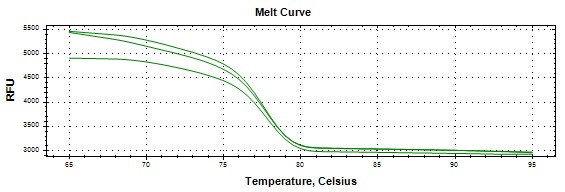

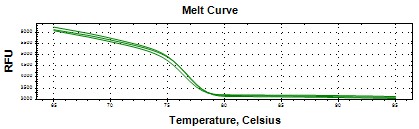

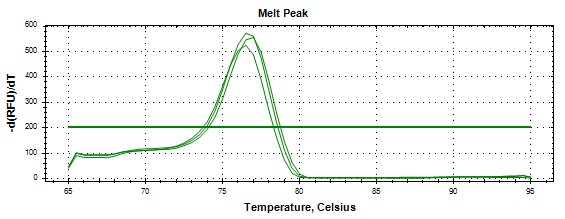

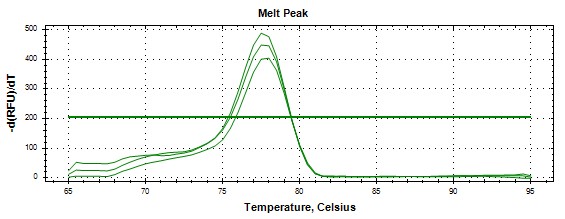

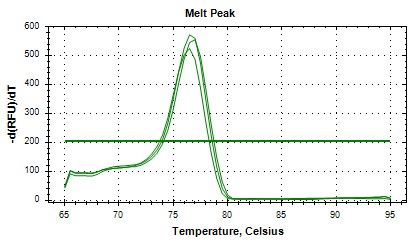

(CP) Tm =77.5	(NP) Tm =78	(TP) Tm =77

*Tight grouping; consistent amplification

**8. *Sr GGDPS***

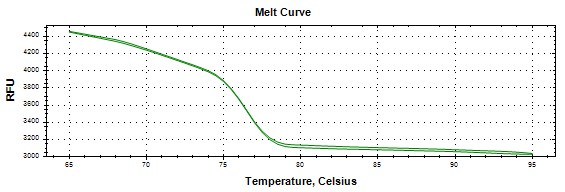

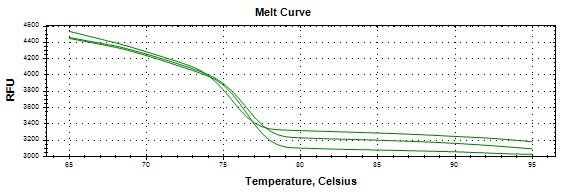

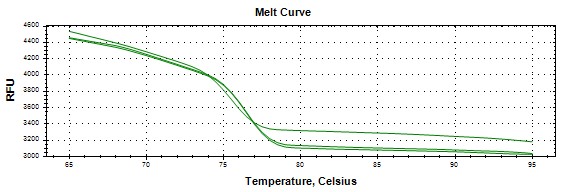

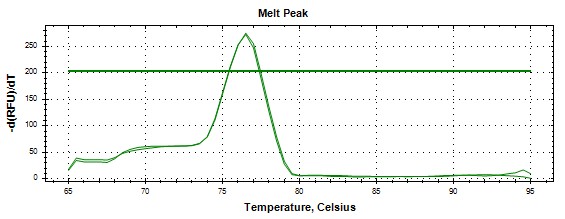

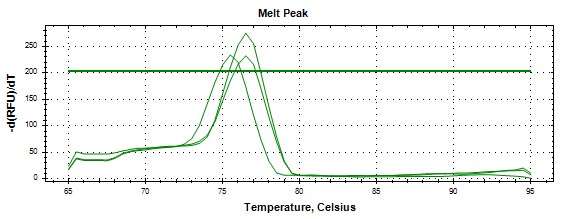

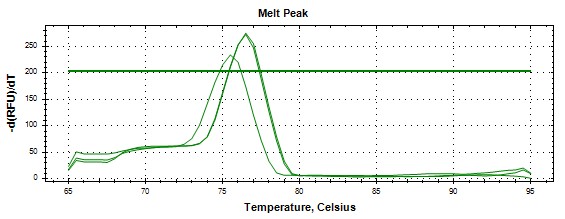

(CP) Tm =76.5	(NP) Tm =75.5	(TP) Tm =76

*Close Tm values; likely specific and stable amplicon

**9. *Sr CDPS***

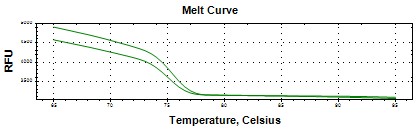

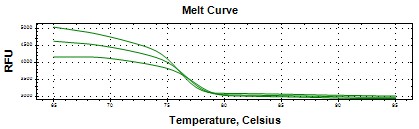

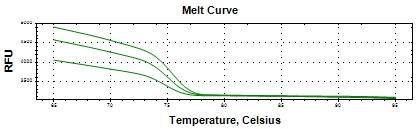

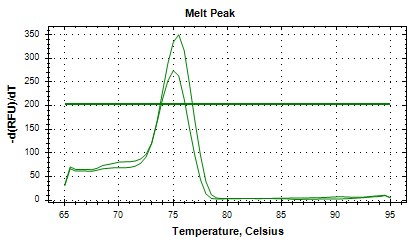

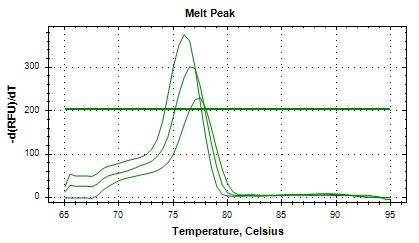

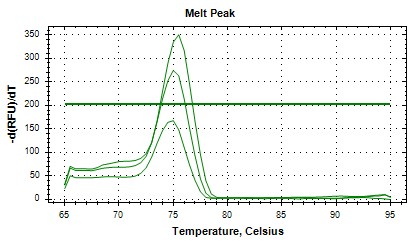

(CP) Tm =75.5	(NP) Tm =76.5	(TP)	Tm =75

*Lower Tm may indicate shorter amplicon

**10. *Sr KO***

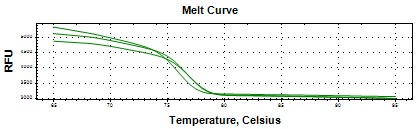

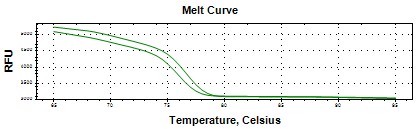

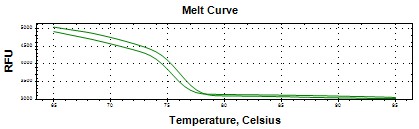

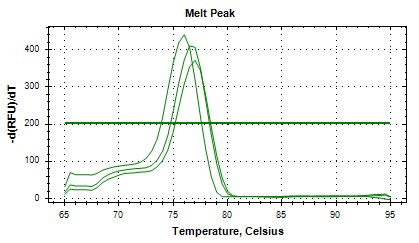

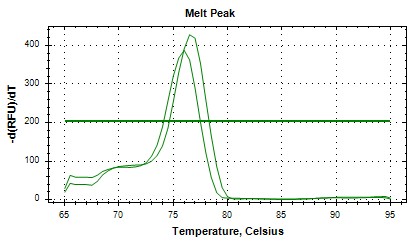

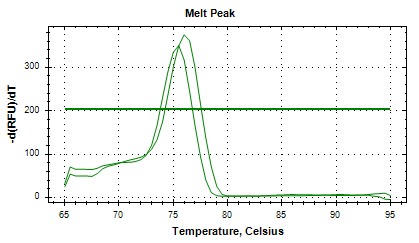

(CP) Tm =76.5	(NP) Tm =75.5	(TP)	Tm =76

*Stable and specific amplification

**11. *Sr KS***

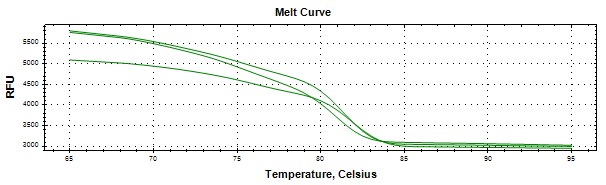

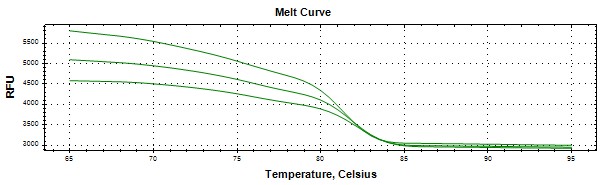

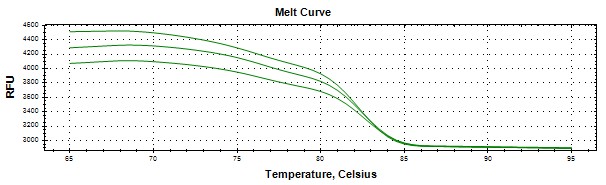

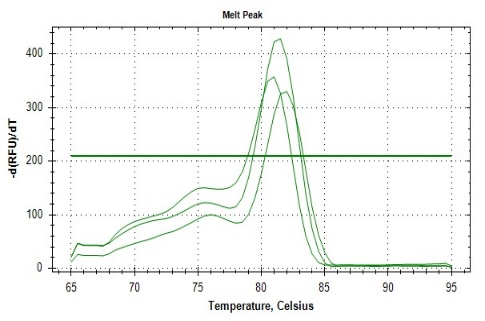

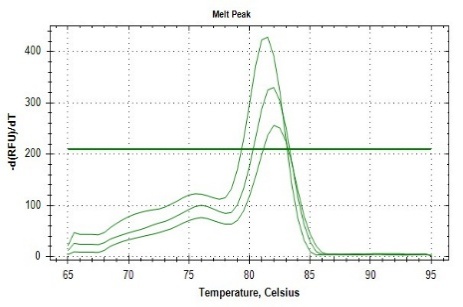

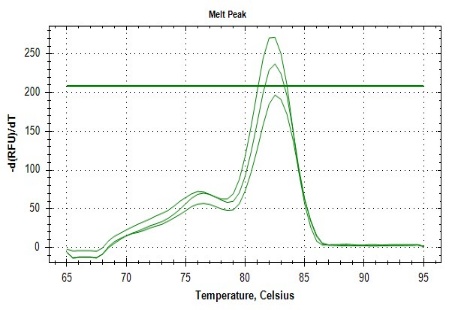

(CP) Tm =86	(NP) Tm =86.5	(TP) Tm =87

*High Tm values, high GC content or long product

**12. *Sr KAH***

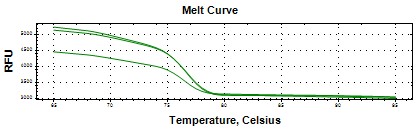

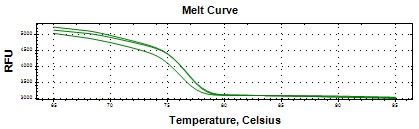

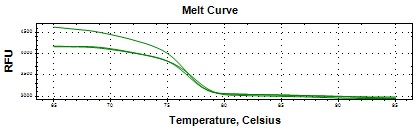

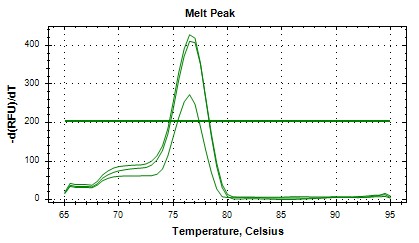

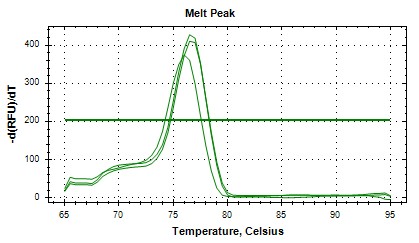

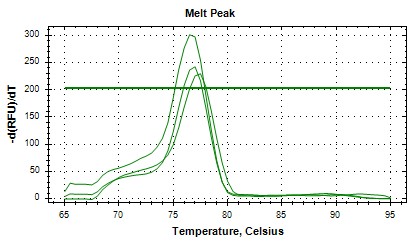

(CP) Tm =76	(NP) Tm =76.5	(TP) Tm =77.5

*Consistent peaks, robust amplification

**13. *Sr UGT85C2***

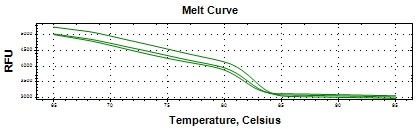

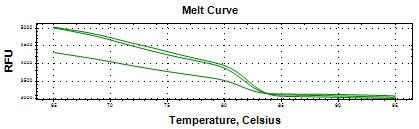

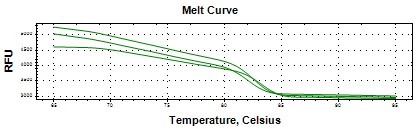

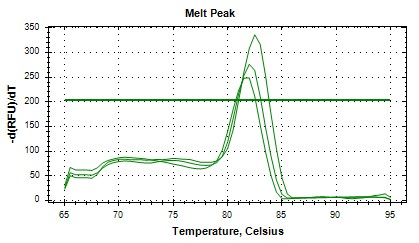

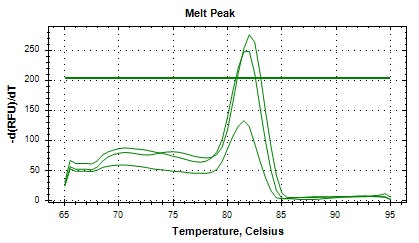

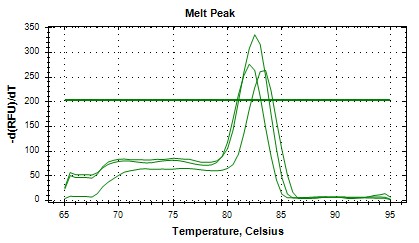

(CP) Tm =83	(NP) Tm =83.5	(TP) Tm =82

*High Tm; complex or GC-rich sequence

**14. *Sr UGT74G1***

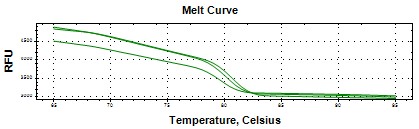

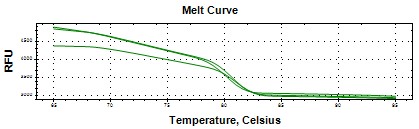

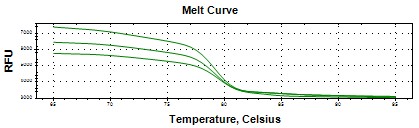

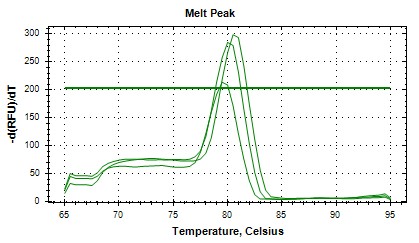

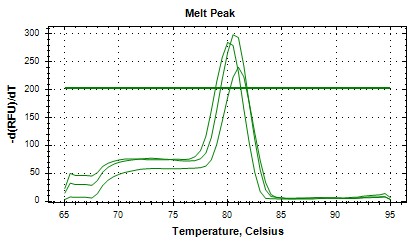

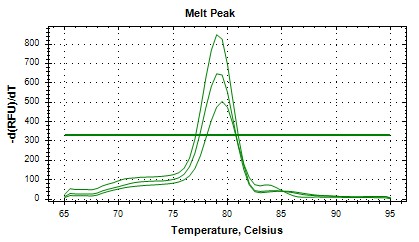

(CP)	Tm =80.5	( NP)	Tm =81.5	(TP)	Tm =79.5

*Tightly grouped peaks, conformational stability

**15. *Sr UGT76G1***

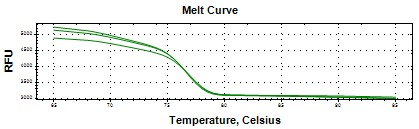

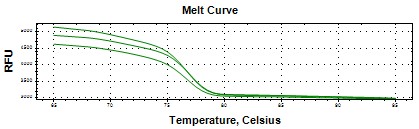

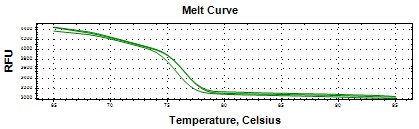

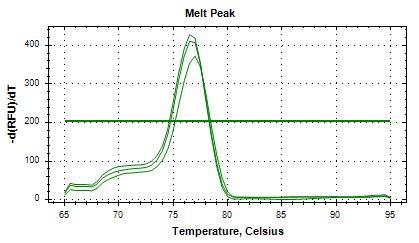

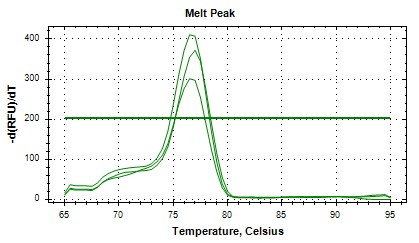

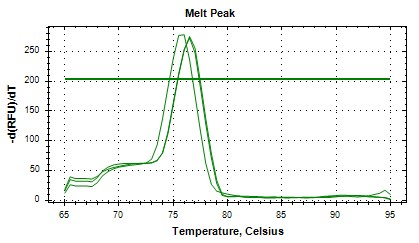

(CP) Tm =76.5	(NP) Tm =77	(TP)	Tm =76

*Specific amplification with low variation

**16. *Sr Actin***

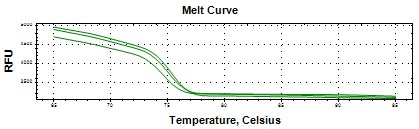

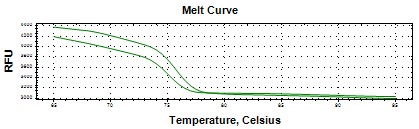

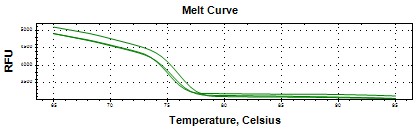

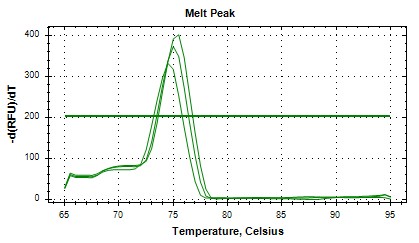

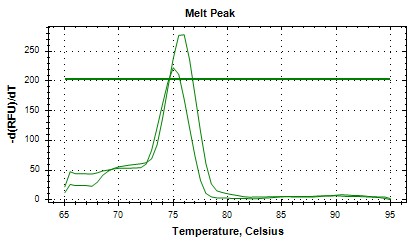

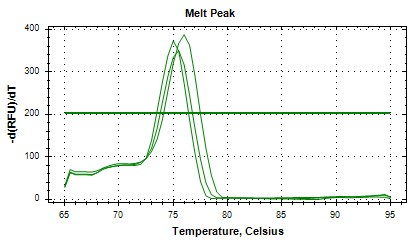

(CP) Tm =75	(NP) Tm =75.5	(TP) Tm =76

*Reference gene with consistent peaks

**17. *Sr GAPDH***

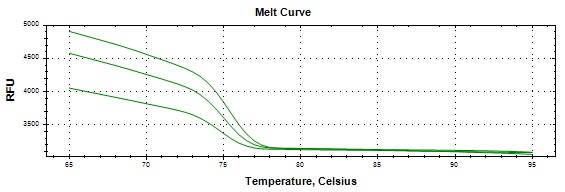

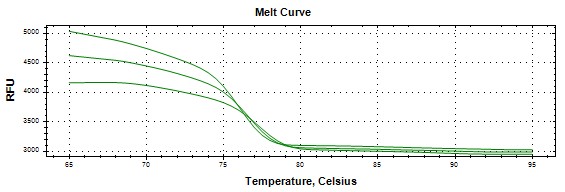

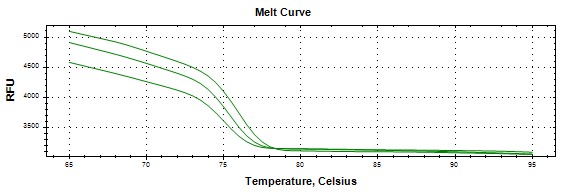

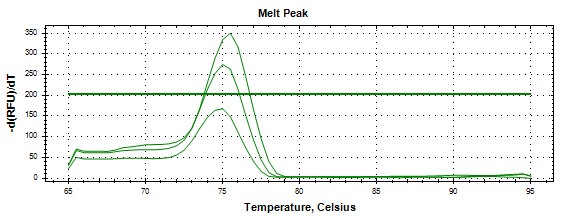

(CP) Tm =75	(NP) Tm =76.5	(TP) Tm =75.5

*Stable housekeeping gene amplification
